# Toothy: an interactive platform for dentate spike curation

**DOI:** 10.1101/2025.10.03.680300

**Authors:** Kathleen N. Esfahany, Amanda L. Schott, Julia M. Grocott, Jordan S. Farrell

## Abstract

Dentate spikes (DSs) are hippocampal population events that occur during low-arousal states, defined by large-amplitude positive voltage peaks recorded in the hilus of the dentate gyrus (DG). DSs can be classified into two types (DS1 and DS2), with DS2 linked to transient increases in arousal, brain-wide neural activation, and functional relevance for memory encoding. Despite growing interest in their physiological and functional properties, no standardized framework exists for detecting and classifying DSs. We present Toothy, an open-source tool for DS detection and classification in large-scale electrophysiological recordings. Toothy offers a modular and interactive workflow comprising three key steps: (1) ingestion and preprocessing of local field potential (LFP) recordings, (2) detection of hippocampal population events, and (3) classification of DS types using peri-event current source density (CSD) profiles. To support both flexibility and reproducibility, features include compatibility with multiple recording formats, customizable processing parameters, interactive event review tools, and comprehensive logging of event detection and classification parameters. In addition to DS detection, Toothy can also detect sharp wave-ripples (SPW-Rs; an oscillatory population event recorded in the CA1 region), enabling comparative analysis between DSs and SPW-Rs. Toothy provides a standardized, reproducible pipeline for DS detection and classification, advancing broader efforts towards investigating hippocampal dynamics across diverse settings.

## Introduction

The hippocampus is a critical structure for encoding, consolidating, and retrieving memories, with much of this processing occurring during offline behavioral states such as rest and sleep^1–4^. During these periods of low arousal, marked by large-amplitude, irregular local field potential (LFP) activity^5^, the hippocampus exhibits two prominent network events: sharp wave-ripples (SPW-Rs) and dentate spikes (DSs). In rodents, SPW-Rs are characterized by high-frequency oscillations (100-180 Hz) in CA1 lasting 20-200 ms^6,7^, whereas DSs are brief (<50 ms), large-amplitude peaks (>1 mV) in the hilus of the dentate gyrus (DG)^8^. Decades of research on SPW-Rs have highlighted their role in supporting memory through replay and consolidation^9–14^, with consensus guidelines recently introduced to standardize their detection^15^. Meanwhile, despite being first characterized around the same time as SPW-Rs, DSs remain relatively understudied. Given the mounting interest in DSs and their role in hippocampal function – including offline/remote memory reactivation^16,17^, neural state switching^18^, and associative memory formation^18,19^ – tools for DS analysis would be especially timely for this emerging field, promoting standardized detection methods and comprehensive reporting of parameters used for defining DSs across studies.

DSs were originally reported to occur at a rate of up to 15 per minute, particularly during immobility and slow wave sleep, and did not occur during locomotion^8^. Since then, reported DS rates have spanned from 2-3 per minute to 2-3 per second, even showing positive modulation by locomotion in some cases^16–22^. This variability in DS rates is likely due to the amplitude-based thresholding for detection, which needs to be chosen carefully to avoid including other positive voltage peaks, such as movement artifacts and gamma oscillations during locomotion. It was also originally reported that DSs come in two types DS1 and DS2^8^ (though see^23^), which later studies demonstrated have strikingly different properties, including local and brain-wide spiking differences as well as coupling of DS2 to arousal^16–18^. Although not universally performed across studies, the gold standard for DS classification is to apply current source density (CSD) analysis^24^ to identify DS1 versus DS2, based on maximal current sinks in the outer or middle molecular layer of the DG, respectively. Precise CSD classification requires the use of linear probes with uniformly spaced electrodes spanning DG molecular layers down to the hilus, although waveform-based classification models, trained on CSD-classified data, can be used with over 80% accuracy in some cases^21^. Given the vast differences in reported rates and differences between DS types, a standardized and comprehensive set of tools for curating DSs would be beneficial to this growing field.

Here we present Toothy, an open-source graphical user interface (GUI) for detecting hippocampal population events (DS1s, DS2s, and SPW-Rs) in large-scale electrophysiological recordings. The GUI provides interactive controls and dynamic visualizations to facilitate the curation of high-quality event datasets. Toothy accommodates diverse experimental contexts and analysis goals by enabling parameter customization, while ensuring reproducibility through automatic documentation of parameter values. The graphical interface makes the workflow accessible to researchers without coding expertise, while the underlying open-source codebase supports bespoke feature development. By consolidating previously fragmented approaches into a single user-friendly platform, Toothy significantly reduces technical barriers to high-quality and reproducible DS detection and classification, thereby opening the doors to new insights into the properties and functional relevance of this understudied hippocampal population event.

## Results

### Overview of Toothy workflow

Toothy is written in Python and utilizes the cross-platform QT framework for its graphical interface, ensuring compatibility with MacOS, Windows, and Linux operating systems. The Toothy software leverages Python’s robust suite of scientific tools to support a modular, interactive pipeline for processing and analyzing multichannel electrophysiological recordings from the hippocampus (**Fig. 1**). Researchers can install Toothy by downloading the underlying code repository from GitHub (https://github.com/Farrell-Laboratory/Toothy/) and following the documentation on the repository page.

**Figure 1.**
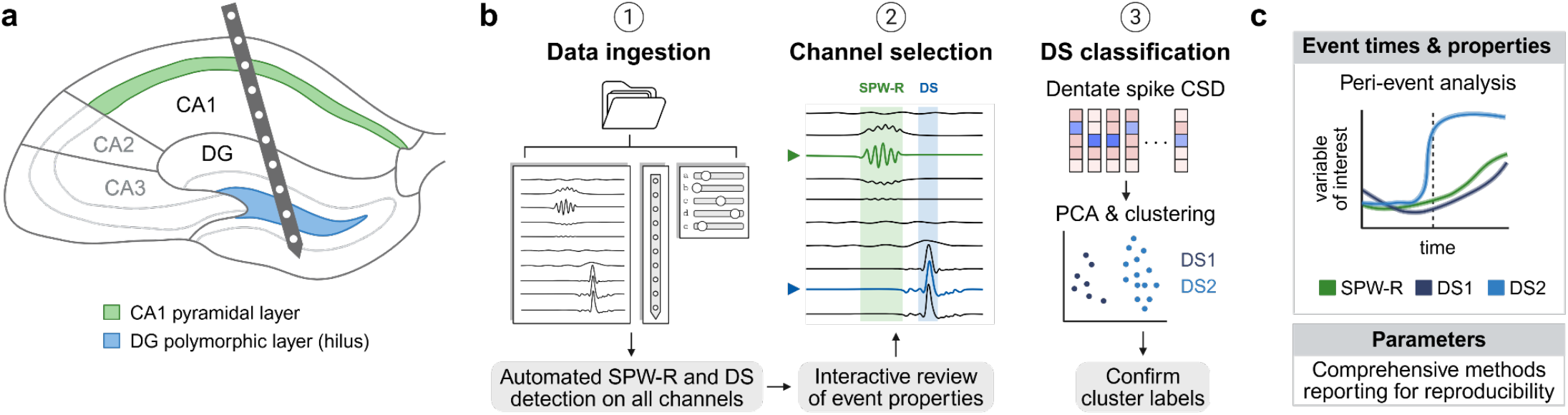
Toothy analysis pipeline for detecting hippocampal population events from electrophysiological recordings. **(a)** Schematic of a laminar probe recording local field potentials (LFP) across hippocampal subregions in the mouse brain, including the CA1 pyramidal layer (green) and dentate gyrus (DG) polymorphic layer or hilus (blue). The multi-channel LFP recording can be analyzed with Toothy to detect sharp wave-ripples (SPW-Rs) and dentate spikes (DSs). **(b)** The Toothy pipeline consists of three main steps: (1) Data ingestion: The user loads LFP data, specifies the associated probe configuration, and sets parameters for further processing. Toothy performs automated event detection across all channels in the recording. (2) Channel selection: For each event type (SPW-Rs and DSs), the user selects a representative channel based on the channel’s properties (selected channel marked by triangle and colored trace; green for SPW-R and blue for DS). Toothy provides interactive features for visualizing and comparing event properties across channels. (3) DS classification: For each DS, Toothy computes a current source density (CSD) profile, finds a 2D representation using PCA, then clusters events into DS1 and DS2. Toothy displays the mean waveforms and CSD profiles for each cluster for the user to use to refine the clustering before confirming the labels for DS1 and DS2. **(c)** Toothy output: Toothy returns a table of event times and properties as well as a comprehensive log of the parameters used for event detection and classification parameters. The event times enable peri-event analysis of other simultaneously-recorded variables of interest, including neural activity (e.g., spiking, photometry) and behavior (e.g. pupil size, whisker movement). The parameter log promotes comprehensive reporting of event detection methods, supporting reproducibility.

Upon running Toothy, the user is greeted with a home screen comprising several buttons, including the main workflow (“Step 1: Load data” and “Step 2: Analyze events”) and convenience menus for functions that can also be achieved within the main workflow (namely setting paths and parameters and creating probe configurations) (**Fig. 2**). The main workflow begins with data ingestion, in which the user selects a recording, specifies the associated probe configuration, and sets parameters for further processing. This step is followed by an internal processing stage in which Toothy scales and optionally downsamples the data, then detects events across all channels. Next, the user enters the main analysis phase of the workflow, channel selection, in which the user selects an optimal channel for identifying DSs or SPW-Rs, facilitated through several visualizations. Finally, the workflow concludes with a DS classification step, in which Toothy performs CSD analysis on the LFP for each DS, performs principal component analysis (PCA) on the CSD, then clusters the CSD to label each DS as type 1 or type 2. Toothy produces two key outputs: (1) a table of event times and properties, which can be used for any number of downstream analyses across concurrent modalities such as imaging and behavior, and (2) a log of the parameters used for each step of analysis, which can be used for methods section reporting to promote reproducibility.

**Figure 2.**
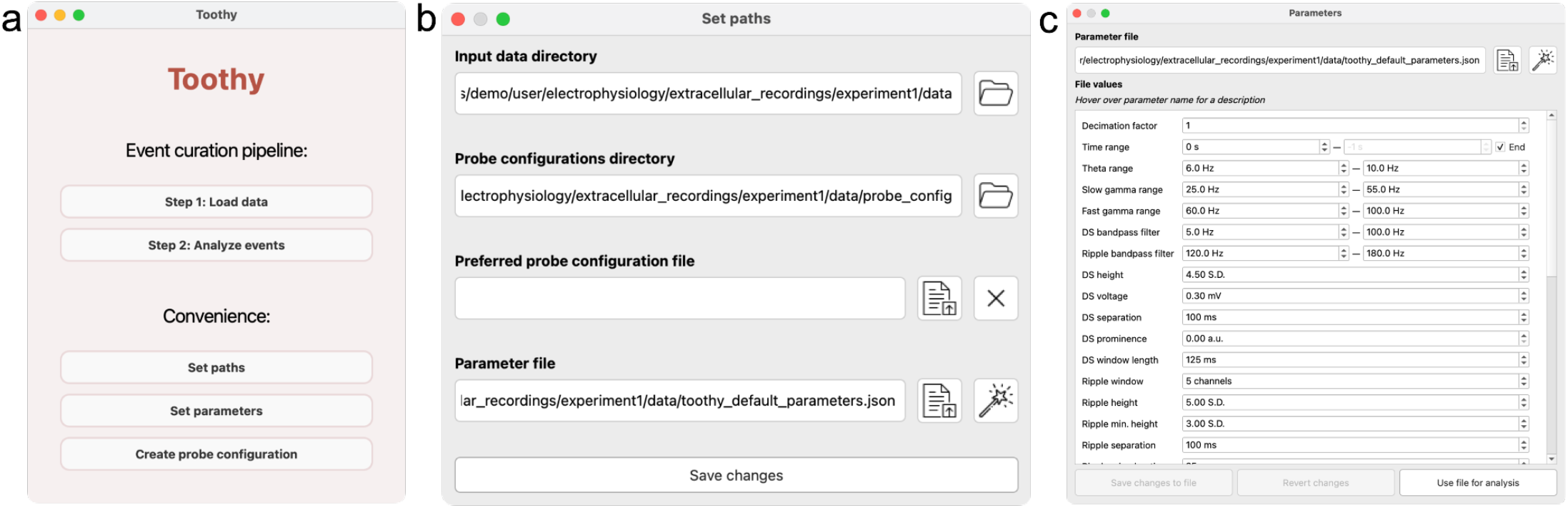
Toothy home screen and convenience windows. **(a) Home screen:** Central access point for all Toothy modules. **(b) Path setting window:** Accessed from the “Set paths” button on the home screen, the path setting window allows the user to set default paths for folders and files to which Toothy will point at later stages in the workflow. **(c) Parameter setting window:** Accessed from the “Set parameters” button on the home screen, the parameter setting window allows the user to choose between using default parameters for signal processing and event detection (optimized to mouse hippocampal data) or using custom parameter files.

### Data Ingestion

The main Toothy workflow begins with data ingestion, which refers to the process of selecting a recording, as well as mapping it to a probe configuration and setting relevant parameters for further processing. These steps can be completed in the Data Ingestion Window of the GUI. To open the Data Ingestion Window, users click “Step 1: Load data” on the Toothy home screen. The Data Ingestion Window proceeds modularly through the following steps: (1) select a recording (**Fig. 3a and 3b**), (2) map the recording to a probe configuration (**Fig. 3c**), and (3) select processing parameters (**Fig 4**). After completing these three steps, users click “Process data” at the bottom of the window to trigger a signal processing pipeline for detecting events across all channels of the recording.

**Figure 3.**
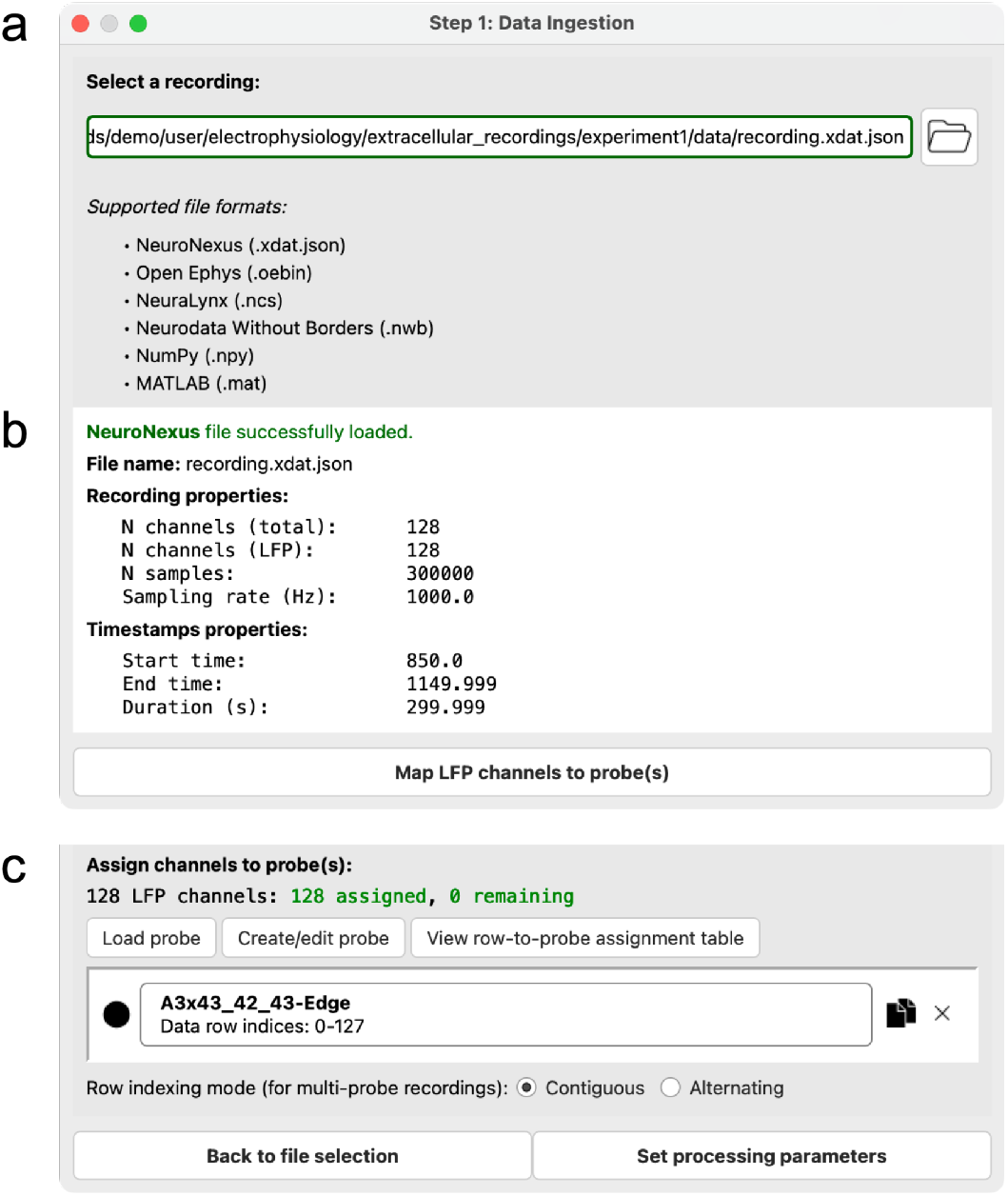
Data Ingestion Window. **(a) Step 1: Select a recording**. Users begin data ingestion by selecting a recording. The user clicks on the folder icon to open a file explorer and select a recording file. Once a file is selected, the box with the file path turns green and the next sections of the window appear, shown in (b), alongside the “Map LFP channels to probe(s)” button. **(b) Metadata from the selected recording**. The properties of the recording (number of channels, number of samples, and sampling rate) and the associated timestamps (start time, end time, duration) are displayed. In this example, a NeuroNexus file with 128 channels sampled at 1000 Hz was loaded. **(c) Step 2: Map the recording to probe file(s)**. A probe file can be loaded or created to map the channels in the file to specific probe(s) and shank(s). Here, a probe configuration corresponding to a NeuroNexus A3x43/42/43-Edge probe. Once all channels have been mapped to a probe, the “Set processing parameters” button is enabled.

**Figure 4.**
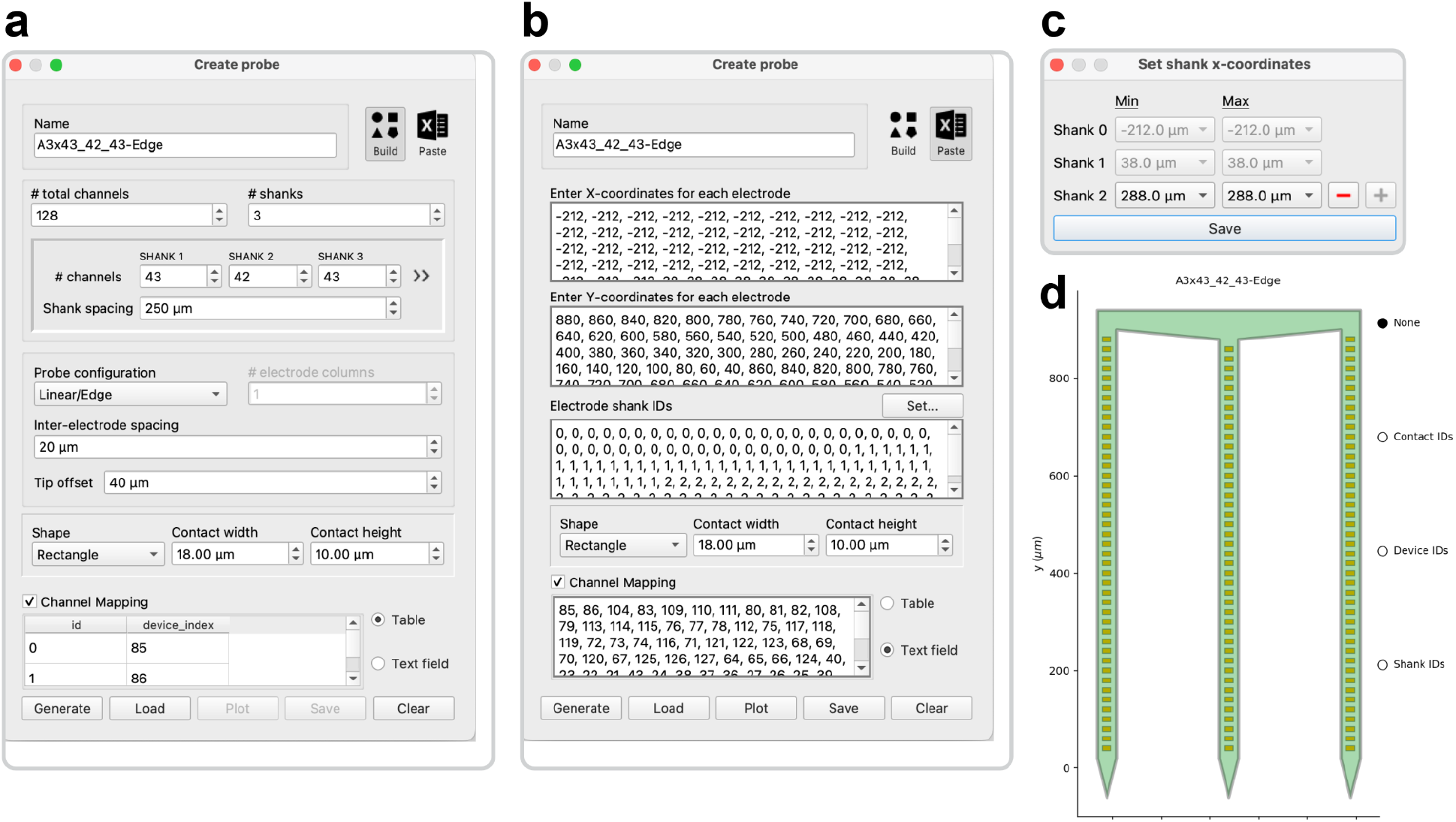
Designing a probe object. **(a)** <u>The Build</u> tool generates a probe object from descriptive parameters, including channel count, shank count, electrode configuration, and contact spacing. Channel mappings can be entered as table rows or comma-separated values to set the probe’s wiring configuration, otherwise the physical probe layout is assumed to match the logical data order. **(b)** <u>The Paste</u> method generates a probe object from arrays of electrode X and Y coordinates. Electrode shank IDs can be entered as a corresponding array, or they can be assigned to specific X-values through a dedicated interface **(c)** This method supports irregular or asymmetrical layouts that violate the assumptions of the “Build” tool. **(d)**The generated probe object is displayed with interactive labeling of contact indices, device indices, and shank IDs. Probe configurations are saved in JSON format for compatibility with the probeinterface package^27^.

To select a recording, users click the folder icon to open a file explorer and select a file (**Fig. 3a**). Toothy supports six file formats, spanning both popular electrophysiology acquisition and storage formats (Neurodata Without Borders, OpenEphys, NeuroNexus, and Neuralynx) and generic array files (NumPy^25^ and MATLAB arrays). Proprietary electrophysiology formats not directly supported can be converted into Neurodata Without Borders (NWB)^26^ files using the [NeuroConv](https://github.com/catalystneuro/neuroconv) Python package or the [NWB GUIDE](https://nwb-guide.readthedocs.io/en/stable/) desktop app before using Toothy. Toothy uses the SpikeInterface Python package^27^ to extract LFP data, timestamps, and relevant metadata (sampling rate and voltage units) from each file. For array files without embedded metadata, Toothy prompts the user to manually enter the information. When a recording is successfully loaded, the window expands to display the properties of the recording (number of channels, number of samples, and sampling rate) and timestamps (start time, end time, and duration) (**Fig. 3b**). Internally, following this step, all recordings are treated identically, regardless of source format.

Next, LFP signals must be mapped to the appropriate recording probe(s). The Data Ingestion Window automatically displays the number of signals detected in the raw data, as well as the number assigned to a valid probe channel (**Fig. 3c**). Users may press the “Load” button to upload a probe object from a saved configuration file. Alternatively, the “Create” button initiates a Probe Designer GUI where users can easily generate a new probe object. The “View” button launches a pop-up table displaying the probe index assigned to each data row, with blank cells indicating unassigned signals. Assigned probes will appear by name in the main window and can be duplicated or deleted using the associated icons. When each data row (i.e. channel) is successfully assigned to a unique probe channel, these items will appear in green and the “Set processing parameters” button at the bottom of the window becomes enabled. Clicking this button initiates Toothy’s internal data processing step, in which LFP signals are filtered and event detection is performed for all channels using previously established methods^18^.

Finally, users confirm the parameters used for processing the recording and performing event detection. Before performing filtering and other signal processing operations to detect events, the data may first be downsampled to reduce the time and storage consumed by Toothy.

### Customizing parameters

Before initiating the data processing step, the user should confirm their chosen analysis parameters used in filtering and event detection. Parameters can be adjusted before starting the Toothy workflow (through the “Set parameters” button on the home screen) (**Fig. 2c**) or from within the Data Ingestion Window. Toothy’s framework for managing numerous configuration parameters allows the user to exert a granular level of control over the inputs to data analyses. The main settings file dictates the values for over fifty parameter inputs that affect all stages of the data pipeline, from signal processing and event detection to CSD estimation and cluster analysis.

Settings are stored in plain text (TXT) files for wide cross-platform compatibility and accessibility outside the GUI environment. Directly editing the parameter values through the text file is possible but not recommended, as this may disrupt the precise file structure required to accurately parse its contents; instead, editing through the GUI is advised. Toothy offers a dedicated interface for viewing and modifying the contents of the current parameter file, accessed by “Set parameters” on the home screen (**Fig. 2c**). The order of parameter rows in the GUI corresponds to the sequential stage of analysis; the first inputs govern raw data processing (e.g. downsampling factor, frequency band cutoffs, and event detection thresholds), followed by inputs for CSD calculation (e.g. estimation method) and finally DS classification (e.g. clustering algorithm). Changes can be saved to the existing settings file or used to create a new one, and the “Reset values” button erases any unsaved changes by reloading the original file.

### Probe configurations

Multi-channel data analysis depends on the unique configuration of the neural probe and its accurate representation in the processing pipeline. While this may be straightforward in a single lab using a few standard probes, it becomes increasingly challenging when analyzing data from varied sources, each with its own probe geometry, size, and channel mapping. A flexible probe designer window allows users to easily create probe configuration files for each of these diverse probe types, enabling rapid customization to match the exact electrode geometry and wiring configuration of any physical probe in use (**Fig. 4**). Toothy uses the probeinterface Python package^28^ to create software representations of probes, stored in a data format based on JSON to ensure compatibility and ease of data sharing across different systems and software environments.

A probe object is defined by its electrode geometry and wiring, edited through two main methods: the Build tool, which generates standard configurations (linear, polytrode, or tetrode) based on user parameters, and the Paste tool, which allows manual entry of arbitrary electrode coordinates for full customization. Additional options include specifying electrode spacing, tip offsets, and contact shapes/sizes, as well as defining channel mappings to link physical electrode contacts with device indices. By default, contact indices follow a consistent ordering scheme, but users can manually override this when hardware rearranges signals.

When all critical parameters for a chosen configuration are set to valid values, the Probe Designer GUI automatically enables the “Generate” button at the bottom of the window. This button synthesizes the current inputs to construct a unique probe object, plotted in a separate window for visual verification of the electrode geometry (**Fig. 4d**). The plot window also allows users to review the probe channel mapping by interactively displaying the contact indices, device indices, or shank IDs above the corresponding electrodes. Once a probe object is generated, the “Save” button allows the data to be written to a file and stored for later use. The default file name uses the text in the “Name” field as a base probe identifier, appends a “_config” tag signifying a probe configuration file, and recommends the `.json` extension for optimal data preservation. The GUI also supports exports to MAT and PRB file formats used by other software packages^29,30^, although some information is lost as a result. A previously saved probe object can be imported by using the “Load” button to select the target configuration file and read in its probe data, which will update the GUI accordingly.

### Event detection

The Toothy data processing pipeline prepares LFP signals for analysis through a series of transformations, manipulations, and preliminary analyses according to the parameters set in the previous phases. The first step is to organize, scale, and downsample the raw data. Using channel mappings from one or more probes in the recording, signals are organized by probe, with each channel array sorted by depth, from shallowest to deepest. Signals are then scaled to standard units of millivolts for consistency. By default, Toothy does not downsample LFP data when the sampling rate is already at 2.5kHz or below. However, to facilitate efficient processing and visualization of data at higher sampling rates, Toothy can downsample data from its original sampling rate (e.g.,kHz, 10 kHz, or 30 kHz) to a user-defined rate (1 kHz by default) by averaging bins of consecutive samples.

Once organized, the LFP signals are bandpass filtered to isolate specific frequency bands with the following default values: theta (6-10 Hz), slow gamma (25-55 Hz), fast gamma (60-100 Hz), ripple (120-180 Hz), and a broader 5-100 Hz band used for DS detection. Each filtered signal is then analyzed for amplitude and calculated as the standard deviation over the entire recording, which provides a mean value for each channel. To facilitate comparisons, these values are normalized across all channels on each probe for visualization the relative power in each frequency band across depth (**Fig. 5d**). Both raw and normalized power values are saved in a summary table for further analysis.

**Figure 5.**
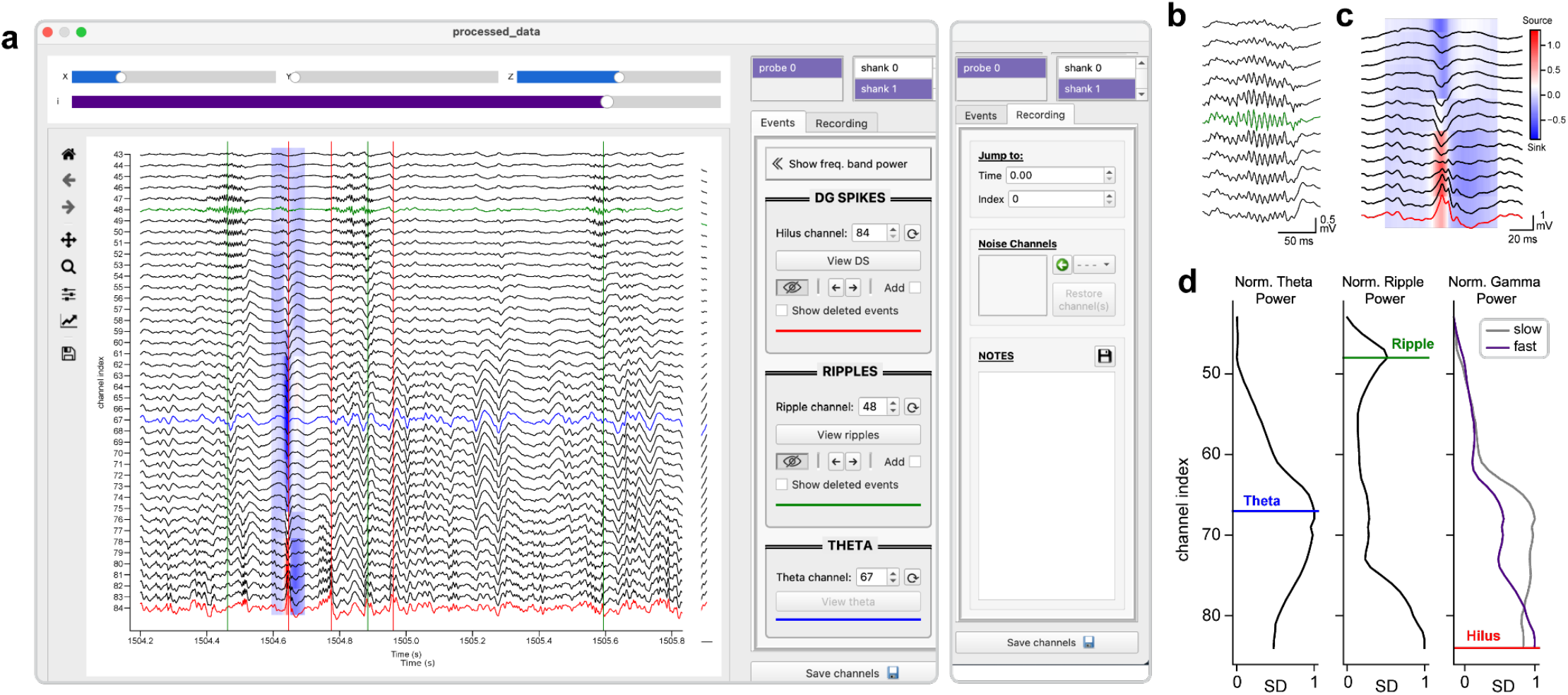
Selecting event channels and curating DS and SPW-R datasets. **(a)** The Channel Selection Window displays raw LFP traces from the selected probe and shank, with the color-coded signals representing the current event channels for DSs (red), SPW-Rs (green), and maximal theta amplitude (blue). The red and green vertical lines mark the timepoints of detected DSs and SPW-Rs, respectively, and the CSD surrounding the first DS event (blue patch) is displayed behind the LFP traces. The Events tab in the sidebar contains tools for setting event channels and analyzing DS and SPW-R datasets, while the Recording tab (right) provides general utilities for navigating, cleaning, and documenting the data. **(b)** Example SPW-R LFPs in CA1, expanded from the upper dashed box in panel **a**. **(c)** Example DS LFPs and CSD in the hilus, expanded from the lower dashed box in panel **a**. **(d)** Normalized amplitude across channels for the theta, ripple, and slow and fast gamma frequency bands, with event channels marked by color-coded lines.

In the next phase, preliminary detection of DSs and SPW-Rs is performed on each channel. For DS detection, positive peaks are identified in the filtered LFP signal using the `find_peaks` function from Python’s SciPy library^31^. Peaks must meet specific criteria: exceeding a user-defined height threshold, having a minimum prominence, and being separated by a minimum time interval to ensure that DSs with multiple peaks are not extracted as discrete events. For the qualifying events, additional waveform properties such as half-width, peak height, and asymmetry are computed using SciPy’s `peak_widths` function within an analysis window around each peak. Since filtered signal peaks may not align precisely with raw signal peaks, each DS is indexed by the maximum raw LFP value within a 20 ms window around the filtered signal peak. This analysis is repeated for all channels, and results are compiled into a single table for storage.

For SPW-R detection, Toothy first calculates the SPW-R envelope by taking the absolute value of the Hilbert transform of the ripple-filtered signal. The instantaneous frequency is then computed by translating the phase from the Hilbert transform across cycles, converting it into frequency values that are clipped to the ripple band (120-180 Hz). SciPy’s `find_peaks` function is again used to detect peaks in the SPW-R envelope which meet the user-defined height and inter-peak interval thresholds. A lower envelope threshold is then applied, and the `peak_widths` function is used to measure SPW-R widths. Additional criteria are used to filter detected SPW-Rs, including minimum duration and mean instantaneous frequency thresholds. Qualified SPW-Rs are indexed by the largest positive cycle within 40 ms of the filtered peak. As with DSs, SPW-R analysis is performed for all channels, and results are consolidated into a table for storage, along with threshold values for each channel.

The final outputs of the data processing phase include the original and filtered LFP signals (scaled and downsampled), frequency band power values, and detailed tables of DS and SPW-R detection results. Data files are organized by probe in a central recording directory, along with probe configuration data and the parameter settings used in processing. This directory serves as a base resource for subsequent analyses and ensures reproducibility across different recordings.

### Channel selection

The channel selection stage of the analysis pipeline is dedicated to refining and validating the detection of DSs and SPW-Rs after automated processing, with the goal of curating high-quality datasets for downstream analysis. Although Toothy detects candidate events across all recorded channels, true physiological events are expected to occur within specific hippocampal subregions, with DSs localizing to the hilus of the DG, due to its unique folded cytoarchitecture^32^. Any detected “events” outside these regions are likely to be spurious or to reflect unrelated activity patterns, such as large gamma oscillations, leaving a narrow range of potential channels for each event type. These channels can be rigorously compared using the Channel Selection Window (**Fig. 5**), which integrates time-series visualization, statistical insights, and interactive tools to identify the optimal event channels and manually edit the data.

The main interface (**Fig. 5a**) consists of a central plot rendered using Python’s Matplotlib library^33^, and a settings sidebar with a suite of interactive elements. In the upper right corner, users can switch between the available probes for the recording, and between the available shanks for the currently selected probe. LFP signals from the specified probe and shank are vertically stacked according to depth and displayed over a given interval, providing an adjustable “viewing window” into the recording. The main horizontal slider (purple) controls the position of the viewing window, enabling users to quickly navigate across the recording. Pressing the left and right arrow keys shifts the window backward or forward by 25% of its time range, providing higher temporal resolution for stepping through local data. The default size of the viewing window is 2 seconds, which can be reduced or expanded by the “X” slider to inspect specific waveforms or evaluate broader patterns. The “Y” slider changes the height of the central figure, effectively stretching the vertical arrangement of LFP channels for enhanced clarity on smaller displays, while the “Z” slider scales the amplitude of individual LFP traces. Another interactive tool is the span selector, a red box dragged horizontally across the plot to visually select a time range. Pressing the “Enter” key calculates an instantaneous CSD for the highlighted interval, temporarily visualized as a colormap overlaying the corresponding LFP signals (**Fig. 5a,c**).

The right-hand sidebar is organized into an “Events” tab with specific tools for analyzing DSs and SPW-Rs, and a “Recording” tab with general utilities. In the latter, the “Time” and “Index” inputs move the viewing window to a given timestamp or recording index, allowing users to efficiently jump between points of interest. Users can also designate individual broken or artifact-contaminated channels as “noise”, excluding them from event detection, CSD computation, and frequency band analyses. Noise channels appear as flat gray traces in the main display, and they can be restored by setting the designation back to “clean”. Finally, the “Notes” section provides built-in documentation through an editable text field, allowing users to record their observations directly within the GUI. This input is linked with a plain text file (`notes.txt`) in the recording folder, enabling preservation of the information across multiple analysis sessions.

The “Events” tab of the settings sidebar is organized into event-specific panels, and editable inputs are used to specify the optimal channels for DS and SPW-R events. A third input specifies the channel with the maximal theta amplitude, used to identify the hippocampal fissure as a landmark for subsequent CSD analysis of DSs (section 5). Initial event channels are roughly estimated based on their spectral properties and identified in the main plot by color-coded LFP signals, using red for the DS (or hilus) channel, green for the SPW-R channel, and blue for the theta (or fissure) channel. Event markers are displayed for DSs and SPW-Rs on the current channels, represented by red and green vertical lines. Any changes to the event channel indexes are reflected in real time by dynamically updating the color-coding and event markers on the main plot, and channel inputs can be reset to their initial values using the adjacent buttons. Toggle buttons in the DS and SPW-R sections allow users to show or hide the corresponding event markers, while pairs of left and right arrow buttons enable direct navigation between detected events. These features are useful for initial event review, facilitating a preliminary understanding of their distribution and waveform characteristics.

At the top of the “Events” tab is a toggle button to show the relative amplitude across theta, ripple, and gamma frequency bands (**Fig. 5d**). Channels on the Y-axes of the frequency band plots are aligned with the main LFP plot for cross-referencing, and the designated event channels are marked by color-coded lines, which are dynamically updated when the channel inputs are modified. These plots compare the current event channels with peaks in the relative amplitude of the corresponding frequency band, providing valuable context for informing channel selections. Together, these visualizations provide a foundation for an informed initial hypothesis of the optimal event channels.

To refine these initial selections, DS- and SPW-R-specific interfaces (**Fig. 6**) are used to compare detailed event properties across channels. The central plot (**Fig. 6a**) is split into two rows, with static quantitative insights on top and interactive waveform visualizations on bottom. The three statistical subplots in the top row summarize key parameters across all channels, including event count and peak amplitude for both DSs and SPW-Rs, with data points color-coded by magnitude for enhanced clarity. The third subplot is event-specific, showing the peak height above the surround for DSs (**Fig. 6a**) or the ratio of ripple to theta power for SPW-Rs (**Fig. 6b**). The distribution of these properties can identify strong candidate channels for each event; in particular, a peak in the ripple-to-theta band ratio and a peak in DS amplitude guides selection of the SPW-R and DS channel, respectively. Data from the currently selected channel can be highlighted with a red outline, using a toggle button in the settings sidebar. In the bottom row, LFP signals are analyzed using two different visualization modes. In “Single” mode (**Fig. 6d,e**), a horizontal slider is used to scroll through individual waveforms in sequence. Users can sort events by attributes such as timestamp, size, and width to view the full range of the distribution and inspect the waveforms at each extreme. In “Average” mode (**Fig. 6a,c**), the mean event waveforms from multiple channels are overlaid, enabling side-by-side comparisons of their morphology. Both modes allow users to adjust the time window, scale the data, and toggle between raw LFPs and the filtered signals used for event detection. Additional options in the settings are used to show and hide various detection thresholds and waveform annotations. These tools balance quantitative metrics with heuristic reasoning to facilitate the selection of optimal event channels.

**Figure 6.**
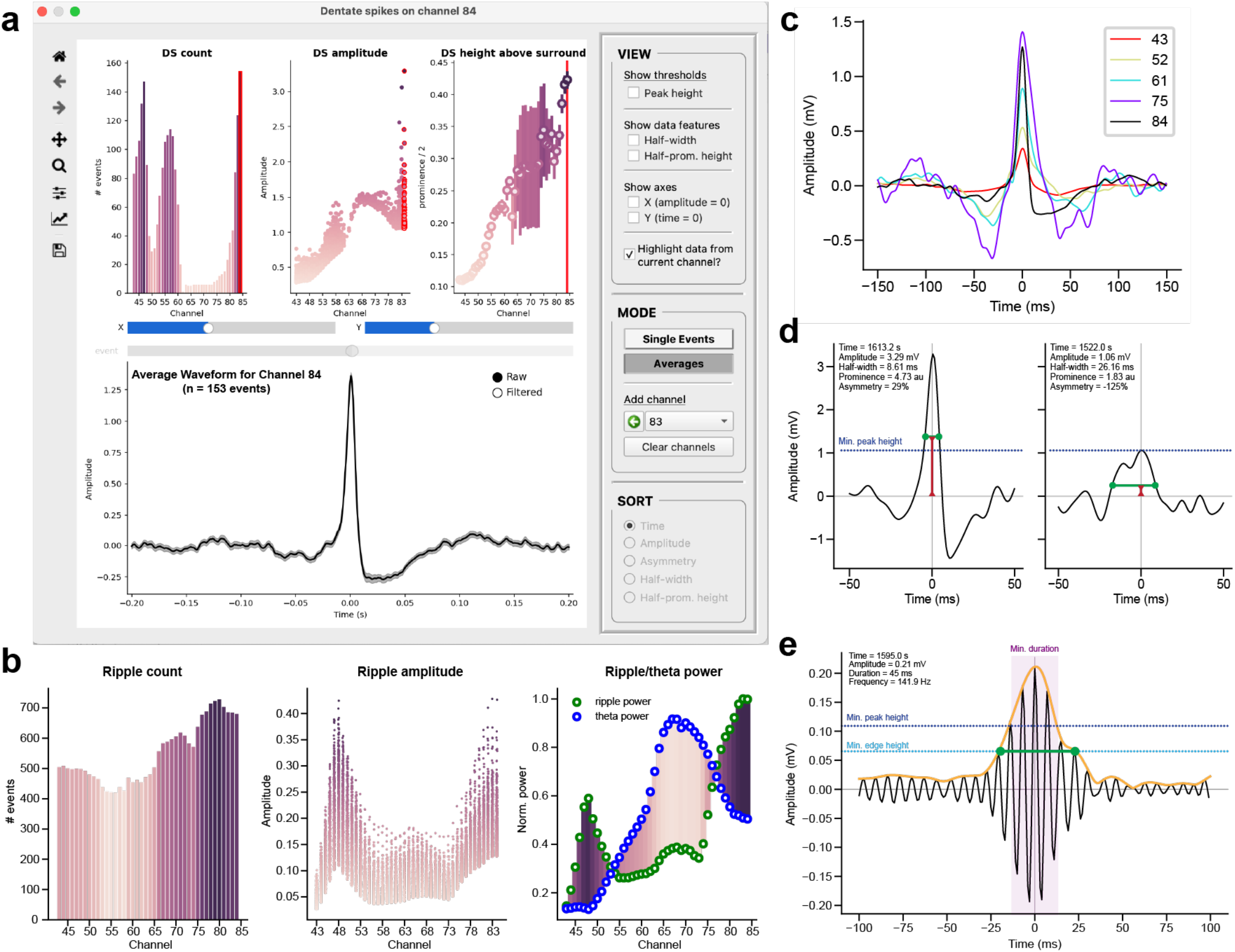
Event-specific analyses. **(a)** Interface for comparing DS and SPW-R events. “Single” mode reviews the individual waveforms in a dataset, and “Average” mode compares the mean event waveforms across multiple channels. **(b)** Parameter distributions for SPW-R events, showing the event count, amplitude, and ripple-to-theta ratio across all channels. **(c)** Mean DS waveforms from five example channels distributed across the probe. The overlaid waveforms highlight laminar differences in polarity, latency, and waveform shape across the hippocampal circuit. **(d)** Individual waveforms from two example DSs with the highest and lowest peak amplitude. The peak detection threshold is indicated by the dashed line, and waveforms are annotated to show with their half-width (green) and half-prominence height (red). **(e)** Example SPW-R waveform and ripple envelope (orange) in the filtered LFP. Detection thresholds are displayed for minimum peak height and edge height (dashed lines) and minimum duration (purple shaded region), and the event duration is indicated by the green bar.

A powerful feature of the Channel Selection Window is the ability to manually curate event datasets, refining the set of automatically detected events by removing artifacts and adding missed events. To add an undetected DS or SPW-R to the dataset, the user must double-click the desired timepoint on the plot while the “Add” option is checked in the corresponding event section. Newly created events are indexed to the LFP peak closest to the clicked point, and they appear as dotted lines for visual differentiation. Users can remove specific DSs or SPW-Rs by dragging the span selector around the desired event markers and pressing the “Backspace” key to temporarily delete the events, or the “Escape” key to permanently erase them. Deleted events can be viewed through a separate toggle button (appearing as dashed lines), and they can be automatically restored by selecting the markers with the span selector and pressing the spacebar. Erased events, however, are permanently removed from the dataset and must be manually re-added by the user. This iterative refinement process generates high-quality, rigorously validated DS and SPW-R datasets for downstream analysis.

The final step of exporting the data to the recording folder is executed using the “Save” button (**Fig. 5**). Event channels are written to an NPY file, while DS and SPW-R datasets are saved as CSV tables. The current probe and shank index is appended to each file name as a unique identifier to avoid confusion in multi-probe and multi-shank recording setups. Successful completion of this step allows access to the DS classification module, the final phase of the Toothy pipeline. While Toothy is primarily geared towards DS curation, users may refer to the previously published consensus statement on methods for further curating SPW-Rs^15^.

### Classification of DS1 and DS2

DSs can be separated into two types based on current sinks in the outer (type 1, DS1) or middle (type 2, DS2) molecular layer of the DG, identified by performing CSD analysis. To classify DS events into DS1 and DS2, Toothy provides a DS Classification Window that streamlines CSD estimation and clustering analyses (**Fig. 7**). The interface offers four modular visualization options: i) an interactive CSD window (**Fig. 7a**), ii) heatmaps of raw and processed signals (**Fig. 7b)**, iii) mean event profiles for DS1 and DS2 (**Fig. 7c**), and iv) clustering results (**Fig. 7d**). The workflow begins by selecting channels for CSD analysis using an interactive slider. By default, the CSD window spans from the selected DS channel (i.e. the hilus) to the selected theta channel (i.e. approximately the hippocampal fissure) to encompass the entire molecular layer of the DG. Users should exclude channels above the fissure from CSD estimation, as the strong sink exhibited by the stratum radiatum during SPW-Rs, which can co-occur with DSs^8,18,22^, introduces an additional dimension into the clustering analysis and may prevent accurate classification of DS types.

**Figure 7.**
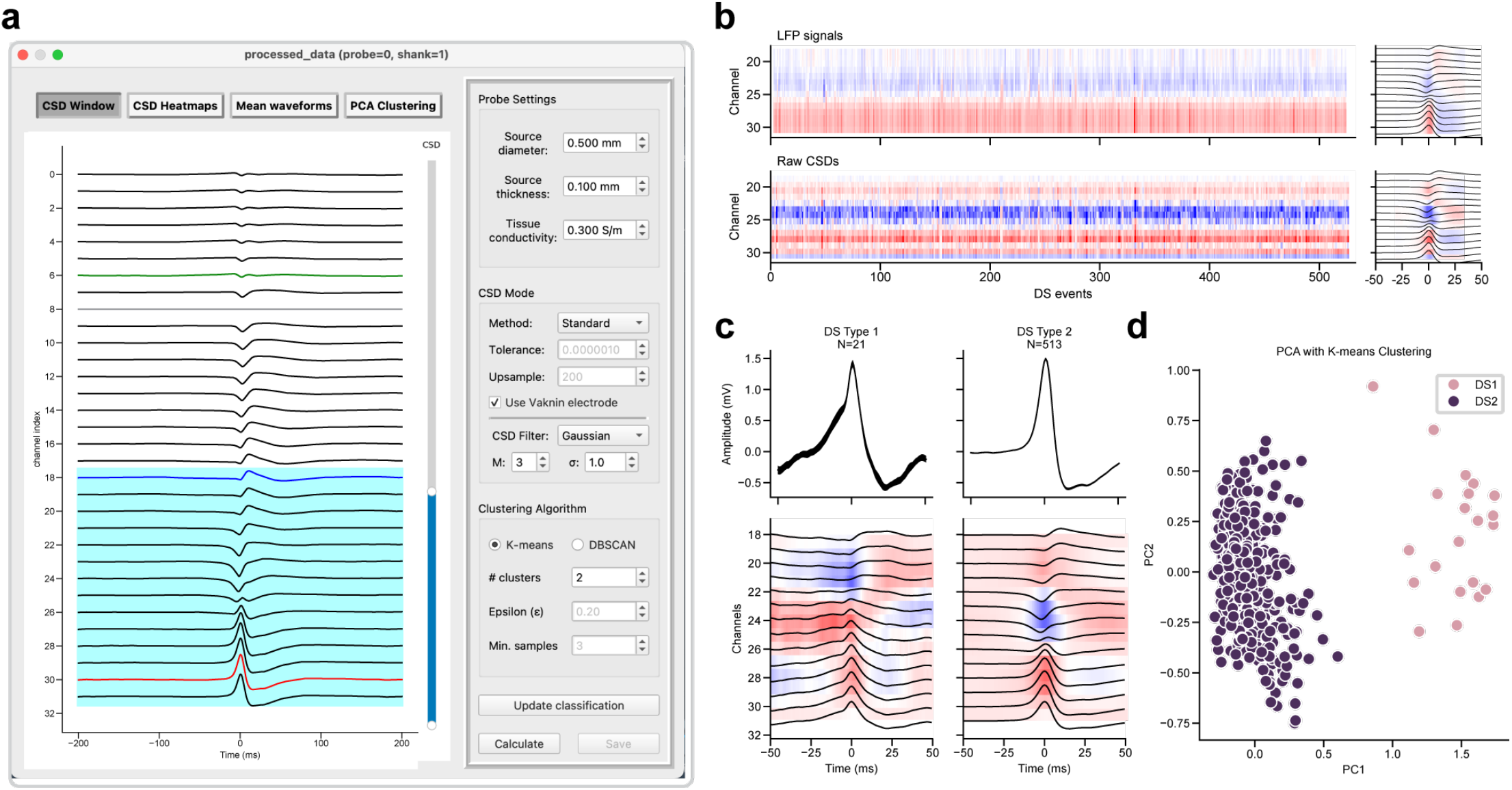
Classification of DS1 and DS2 using CSD estimation and event clustering. **(a)** Interface for CSD-based classification. The interactive CSD window is used to define the range of channels for analysis, and the sidebar enables configuration of CSD calculation settings and clustering algorithm parameters. The “Calculate” button initiates the CSD estimation and PCA-based clustering, and the results are displayed in three additional visualizations accessible through the inputs above the plot. **(b)** Heatmaps of raw LFPs (top) and raw CSDs (bottom) for each DS event. Right, mean LFPs and CSD surrounding DSs. **(c)** Mean LFP waveforms and filtered CSDs for DS1 and DS2 events. **(d)** Two-dimensional PCA projection of the normalized filtered CSDs. Events are color-coded by cluster label (DS1 or DS2).

Users may choose the standard CSD method or one of several inverse CSD (iCSD) approaches, which include δ-source, step, and spline methods^24^. By default, Toothy uses the standard CSD method with Vaknin’s procedure and third-order Gaussian smoothing (σ=1), but alternative CSD methods will have a negligible impact on the subsequent clustering. Heatmaps are used to display the LFPs and the computed CSDs (**Fig. 7b**). Summary plots display the averaged LFPs and CSDs within a 100 ms window surrounding each DS, with the color gradient indicating the signal intensity across channels and events.

Following CSD estimation, principal components analysis (PCA) is performed on the normalized filtered CSD profiles. Under physiological conditions, the PCA-based analysis should produce two distinct clusters, corresponding to DS1 and DS2 events, as evident on the PCA scatterplot (**Fig. 7d**). Points are color-coded by classification as DS1 or DS2, based on the clustering results from K-means or DBSCAN algorithms, available for comparison via toggle buttons on the plot. K-means requires a predefined number of target clusters, while DBSCAN requires parameters for epsilon (the local radius for expanding clusters) and the minimum number of samples per cluster (note DBSCAN can produce a third “undefined” cluster). Toothy automatically labels the clusters by comparing their mean CSDs, designating the cluster with a maximal sink closer to the fissure as DS1 and that with a sink closer to the granule cell layer as DS2. The mean LFP waveforms and associated peri-event CSDs are displayed for DS1 and DS2, along with the number of events in each category (**Fig. 7c**). These plots allow users to compare the classification results with expected physiological patterns, and users can switch the label assigned to each cluster if visual inspection determines that this automatic labeling was inaccurate. Following successful classification, the user can press “Save” to add the event types to the DS dataset. This step concludes the Toothy workflow, producing a set of CSV files with event times and types that facilitate any number of downstream analyses.

## Discussion

Toothy addresses a critical gap in the field of hippocampal dynamics: the lack of standardized and accessible methods for dentate spike detection and classification. By providing a user-friendly, open-source platform for curating high-quality datasets of hippocampal events, Toothy opens the door to a wide range of downstream analyses to investigate the properties and functional relevance of DSs. The modular, customizable pipeline – encompassing data ingestion, channel selection, and DS classification – promotes flexibility across experimental contexts while ensuring reproducibility through automatic parameter logging.

Toothy functions not as a black-box detector but as an interactive tool that supports user-guided curation. While Toothy reduces technical barriers by consolidating complex analytical steps into a graphical interface, effective use still requires familiarity with hippocampal LFP events. For example, an ideal user would know to consider the theta/ripple band ratio in selecting a SPW-R channel, the DS height and mean waveform in selecting a DS channel, and the relative sink position in confirming the automatically-assigned DS1/DS2 classification labels. While Toothy selects event channels and classification labels using heuristics developed primarily from dorsal hippocampal recordings in mice, the unique nature of each recording necessitates a user with some prior knowledge of hippocampal events to ensure proper completion of each step. In addition to user expertise, Toothy’s performance depends on the quality and properties of the input data. The curation of appropriately separated DS types requires sufficient channels in the molecular layer and hilus and sufficient sampling of quiescent behavioral states in which DSs occur (awake rest and non-REM sleep). Nevertheless, given that Toothy’s clustering of event types uses PCA, which is agnostic to the location of current sinks and sources, irregular sampling of DG LFP with alternative electrode configurations (e.g., wire bundles or tetrodes) may still yield distinct DS clusters that can be appropriately classified based on waveform differences^21^ and behavioral features^18^.

Ultimately, Toothy provides a foundation for reproducible analyses of dentate spikes. Standardized detection and classification of DSs and SPW-Rs will facilitate comparisons across studies and enable integration with behavioral or imaging data.

## Acknowledgements

K.N.E. is supported by an NSF GRFP award. A.L.S. is supported by NIH NINDS T32 training program (T32NS007473). Dentate spike research in the laboratory of J.S.F. is supported by NIH NINDS (R00NS126725), Esther A. & Joseph Klingenstein Fund, Simons Foundation, CureSHANK, CURE Epilepsy, and the Harvard Brain Science Initiative Bipolar Disorder Seed Grant.

